# Improving a removal model for evaluating density changes in a widespread invasive species

**DOI:** 10.1101/2025.10.16.680799

**Authors:** John R. Foster, Kim M. Pepin, Maxwell B. Joseph, Michael A. Tabak, Ryan S. Miller

## Abstract

Invasive species pose a significant threat to native ecosystems. Effective management of invasive species requires accurate abundance estimates through time, particularly at large spatial scales. Traditional methods of abundance estimation from management data, such as camera trap studies and harvest data, face challenges due to variations in sampling protocols and effort standardization. We propose an extension to a previously calibrated removal model by integrating into an Integrated Population Model (IPM) and incorporating multiple removal techniques. The model utilizes management and land cover data, both of which are readily available in many invasive species systems. We evaluate the model with a simulation study, then apply it to management data on wild pigs (*Sus scrofa*). Simulations showed that the model performs well across a wide range of densities and population trajectories. Density estimates were the most biased when effort and density were low and sharpshooting or fixed wing aircraft were used to conduct removals. We discuss the advantages and limitations of using removal models for invasive species management, how the model performs with respect to previous removal modeling efforts, how performance might be improved through changes to data management practices, and the model’s potential to inform decision-making processes for invasive species management.

**Data availability:** The data for this manuscript is available from Dryad (http://datadryad.org/share/gFz3ClanB-8yPB5lQ1x-dl1JqI07CP3SX92P-9UyfaU). The code for this manuscript will is available from Zenodo (10.5281/zenodo.15784366).

## 1 Introduction

Abundance estimation is a foundational element of wildlife management and control of widespread species (Gerber & Kendall, 2018; Nichols & Williams, 2006). However, predicting the spatio-temporal trends in abundance, especially over large areas, can be expensive and challenging (Tingley et al., 2015; Williams et al., 2002) due to computational requirements, sampling biases, complex data governance and the need to integrate disparate data (Sólymos et al., 2013; Zipkin et al., 2021).

That said, national and continental scale abundance estimates are routinely generated for waterfowl and fisheries for population management. For example, abundance estimates for the eastern mallard (*Anas platyrhynchos*) across the Atlantic Flyway integrated data from harvest surveys and mark-recovery data collected by both United States and Canadian governments (Roberts et al., 2023).

While state and/or local scale data are frequently used to estimate abundance of terrestrial species, this is not commonly done at the national scale for game species. The lack of standardized effort and data collection bias can lead to inaccurate abundance estimates when applying population models to hunting data (Lukacs et al., 2011; Aubry et al., 2020). This may explain why there is no national abundance model for the most studied and harvested game species in the United States (U.S.), the white-tailed deer (*Odocoileus virginianus*) (Readyhough et al., 2024).

Invasive species are a special case of game species, where abundance estimates at national scales are needed, but instead of managing populations for social good (e.g. sustained harvest), the goal is to eradicate them. Invasive species outcompete native species and lack population controls leading to uncontrolled growth (Crooks, 2002; McDonough et al., 2022) and devastating consequences at multiple scales (Banks et al., 2015; Hobbs & Humphries, 1995; Mack et al., 2000; Simberloff, 2011). Such spread can threaten native ecosystems, biodiversity, and the economy (McClure et al., 2018; Olson, 2006; Pimentel et al., 2005; Walsh et al., 2016). At broad scales, elimination efforts must be adaptive, requiring sustained funding, efficient management, and coordinated efforts among multiple jurisdictions (Bryce et al., 2011; Epanchin-Niell & Wilen, 2015).

In invasive species management, removal models can inform decision-making processes and evaluate control strategies (Barron et al., 2011; Davis et al., 2016, 2017, 2022; Ramsey et al., 2009; St. Clair et al., 2013). Removal models estimate abundance by analyzing the relationship between the number of animals removed from the population in a given time period, capture probability, and effort (Skalski et al., 2005a), where removals and effort must be reliably recorded, preferably during regular and controlled activities (Skalski et al., 2005b). These models can be used to estimate abundance if multiple removal methods are used (Maunder & Punt, 2004; McCluskey & Lewison, 2008; Rivera & McCrea, 2021). They allow for simulation of different management scenarios to assess the potential effectiveness of removal activities, enabling managers to identify the most efficient and cost-effective strategies for reducing invasive species populations (Davis et al., 2022; Snow et al., 2024). However, removal models can be biased when density is low (Keiter et al., 2017).

The effectiveness of control measures vary spatially and temporally at the national scale (Pepin et al., 2019), which complicates model inference and management (Glow et al., 2020). The limited availability of data in space and time can constrain the development and validation of removal models for application at broad spatial scales. For example, actions such as trapping, sharp-shooting, and aerial operations may not always achieve the intended results due to imperfect detection, incomplete removal, behavioral responses to removal activities, or alteration of population dynamics (Davis et al., 2018, 2022; Tabak et al., 2018). This can lead to biased or incomplete datasets, compromising the accuracy and reliability of model predictions (Keiter et al., 2017).

We aim to overcome these challenges by extending a recently developed removal model for wild pigs (*Sus scrofa*) (Davis et al., 2022) by adapting it into an Integrated Population Model (IPM) framework. IPMs can compensate for data deficiencies by combining locally sparse data from multiple study sites or sub-populations (Schaub & Abadi, 2011). This allows for the estimation of regional or metapopulation trajectories and population demographics, overcoming limitations of independent analyses and accounting for multiple sources of uncertainty due to data limitations or environmental variation (Saunders et al., 2019).

We model survival and fecundity as mechanisms for population regulation (Fig. 1) and expand the data generating process to include the most common removal methods for wild pigs (sharp-shooting, trapping, snaring, and aerial operations) by estimating method-specific capture rates as a function of standardized effort, land cover, and search area.

**Figure 1.**
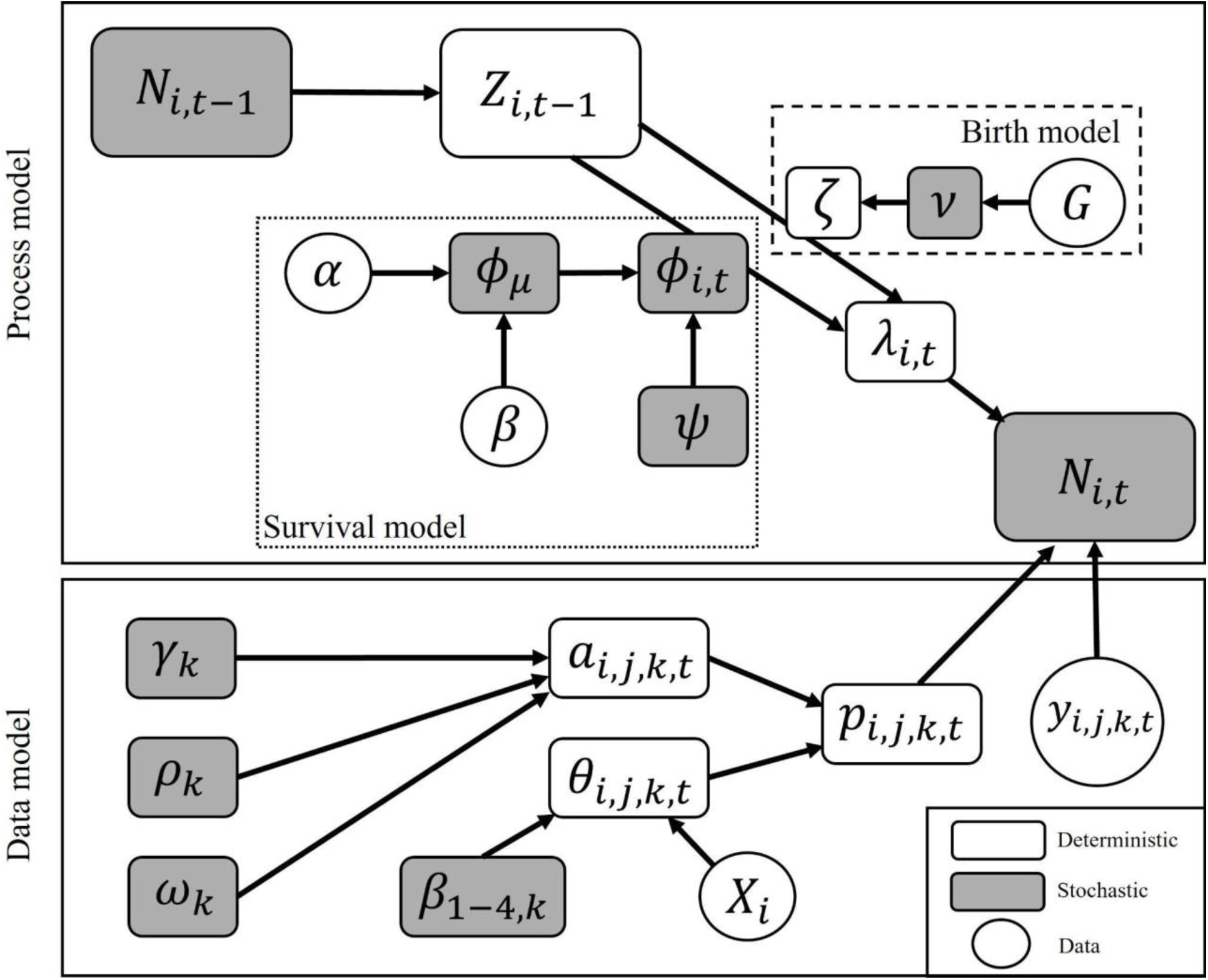
Directed acyclic graph of the removal integrated population model. The subscripts represent property (*i*), removal pass (*j*), removal method (*k*), and primary period (*t*). Circles represent data and model inputs, grey squares represent stochastic nodes, and white squares represent deterministic nodes. Within each property, abundance (*N*) is a function of the survival (dotted box) and birth (dashed box) processes operating on the number of individuals remaining after removals in the previous timestep (*Z*). Hyperparameters *α* and *β* for the global mean survival rate (*ϕ*_*μ*_) were calculated as shown in Supplementary Information 1.1.

A simulation experiment is used to validate the model and quantify its performance at different density levels. Then we apply the removal IPM to estimate the abundance of wild pigs over different ecological contexts in the southeastern U.S. using management data. Our main objectives are to (1) develop a model using management data that is applicable in different ecological contexts, (2) evaluate the data needs and limitations of this model using simulations, and (3) challenge the model to evaluate its robustness in practice.

## 2 Methods

### 2.1 MIS data

Wild pig removal data was obtained from the Management Information System (MIS) of United States Department of Agriculture (USDA) Wildlife Services. We extracted data from 2014-01-01 to 2024-06-30 for the continental U.S. (Table S1), the fiscal year during which the Animal and Plant Health Inspection Service’s (APHIS of USDA) National Feral Swine Damage Management Program began. Data before this time frame were inconsistent. MIS maintains information at the property scale and includes property size (i.e. area), dates of removal activity, removal method(s), units deployed (e.g. number of traps), effort (Table 1), and number of pigs removed for each removal event.

**Table 1.** Parameters used to simulate each removal method. The effort column describes how effort for each method was measured/calculated in the MIS data. The right three columns describe the distributions (U = uniform, G = gamma) from which parameters in the data model were randomly chosen during the simulation experiment. *ω* is the proportion of unique area searched when additional snares and traps are deployed, *ρ* is the scaling factor between effort and area (km^2^) searched by each method, and *γ* is the saturating constant for snares and traps. Empty values indicate where parameters were not used for specific methods. Hobbs hours are the length of time the motor is running (includes flight time to and from the property).

| Method | Effort | $\omega$ | $\rho$ | $\gamma$ |
| --- | --- | --- | --- | --- |
| Sharp-shooting | The number of hours spent and number of people engaged in sharp-shooting | | $U(0.01, 5)$ | |
| Fixed Wing | Hours flown in Hobbs hours | | $U(1, 200)$ | |
| Helicopter | Hours flown in Hobbs hours | | $U(1, 200)$ | |
| Snares | Trap nights | $U(0, 1)$ | $U(0.1, 15)$ | $G(7.7, 4.4)$ |
| Traps | Trap nights | $U(0, 1)$ | $U(0.1, 15)$ | $G(3.6, 3.5)$ |

The data does not include sex, age, location (i.e. longitude/latitude), or any other identifying characteristics of the individuals removed. The finest resolution spatial data was a property identifier that could be linked to a county but not the location of the property within the county. Properties less than the median estimated home range size of pigs (1.8 km^2^ from Kay et al. (2017)) were removed because we could not estimate movement of pigs across properties from our data.

### 2.2 Integrated Population Model (IPM)

Wild pig density was estimated with a state-space integrated population model. The ecological process estimates population growth between 28-day primary periods. A primary period is the length of time that removal events are binned into with the assumption that the population is closed during the interval (Link et al., 2018).

Populations were defined at the property level and partitioned into two processes. The first, defined as *survival* (Supplementary Information 1.1), is the rate at which pigs remain in the population, *ϕ*_*i*,*t*_ (i.e. does not die and does not emigrate). The second, defined as *births* (Supplementary Information 1.2), is the rate at which pigs are added to the population, 𝜁 (i.e. per-capita reproduction and immigration). These processes occur between primary periods, satisfying the closure assumption within a primary period.

#### 2.2.1 Process model

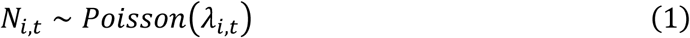

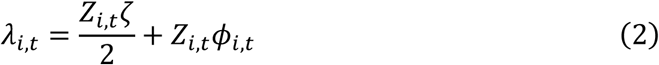

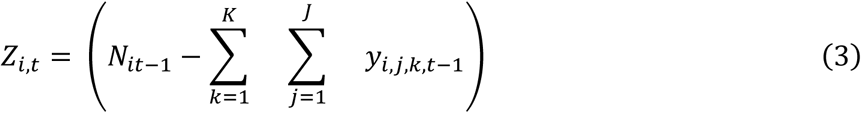

The expected abundance of pigs, *λ*_*i*,*t*_, in property *i* at primary period *t* is the sum of survival and births divided by two assuming the population has equal sex ratios, which the empirical data suggests is a valid assumption (Snow et al., 2020). *Z*_*i*,*t*_ is the number of pigs remaining after removals using *K* methods over *J* passes. Demographic stochasticity was added giving the true abundance of pigs, *N*_*it*_.

#### 2.2.2 Data model

We divided the data generating process into two categories: active and passive. The active methods (sharpshooting, fixed wing aircraft, and helicopters) describe removal events where units moved across the landscape. The passive methods (traps and snares) describe removal events where units were placed on the landscape and left to capture individuals if individuals happened to come across them.

For active methods, we assumed that the search area scales linearly with effort

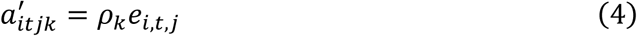

for each removal event where *ρ*_*k*_ is the linear scaling factor on effort per unit deployed, 𝑒_*i*,*t*,*j*_. We assumed the active methods moved at a constant pace, so that one hour of searching increased the area searched by the same amount.

The passive methods were modeled as

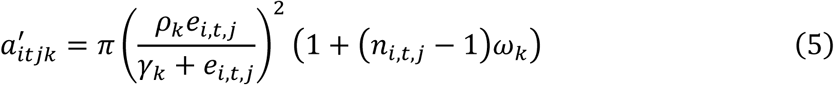

where *γ*_*k*_ is a saturating constant. The different representations of the search area between active and passive methods were because the passive methods did not “wander” when deployed for a longer amount of time. Additionally, we accounted for potential overlap in sampling areas for snares and traps by assuming each unit contributed equally to total effort and defining *ω*_*k*_ as the proportion of unique areas sampled. This was independent of property size and describes the average amount of overlap in search area when 𝑛 traps or snares were deployed. Finally, search area was modified to not exceed the area of the property (𝑎_*i*_) using 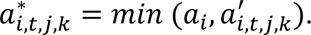

Landscape features can affect the effectiveness of removals (Davis et al., 2018). We expanded on this to differentiate the effect of landscape features by removal method. For example, tree canopy cover may affect a helicopter’s ability to locate wild pigs because they would be harder to identify through dense foliage compared to a trap that was placed in dense canopy cover where pigs are known to inhabit (Lewis et al., 2017; McClure et al., 2015).

The removal rate was a function of landscape covariates, 𝑋_*i*_, and *β*_*k*_ are method-specific coefficients. This was scaled to the proportion of the property searched. The removal probabilities for all passes in a primary period were reparametrized from a multinomial likelihood to a Poisson likelihood (Dorazio et al., 2005; equation 7), and used to determine the number of pigs removed, 𝑦_*i*,*t*,*j*_.

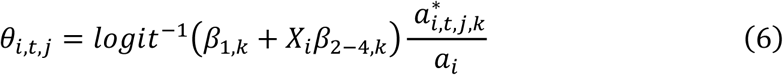

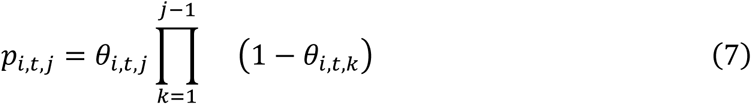

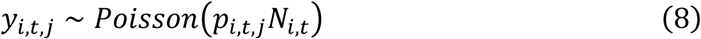

Three landscape characteristics were used to represent potential limitations on detection and capture of wild pigs - percent tree canopy cover, road density, and terrain ruggedness. The moderate-resolution imaging spectroradiometer (MODIS) vegetation continuous field data was used to describe percent tree cover at a 250-meter resolution (DiMiceli et al., 2015). The average percent tree cover was calculated for each county. Terrain ruggedness was calculated using Riley et al. (1999) terrain ruggedness index using 30-meter resolution digital elevation models collected by the Shuttle Radar Topography Mission (SRTM) (Jpl, 2013). Calculation of terrain ruggedness was done using the R spatialEco package (Evans & Murphy, 2023). The average terrain ruggedness index was calculated for each county. Road density was calculated using U.S. Census Bureau TIGER Line data that contains all known U.S. road infrastructure. TIGER road infrastructure data was acquired using the R tigris package (Walker, 2016). Road density was calculated by summing the total linear length of all roads (km) in a county and dividing by the area of the county. Percent canopy cover, road density, and average terrain ruggedness were centered and scaled to one standard deviation.

### 2.3 Simulation experiment

The purpose of the simulation experiment was (1) to evaluate the ability of our IPM to recover known parameters and abundances, and (2) to elucidate data characteristics that lead to accurate abundance estimates. Therefore, we constructed a set of simulated properties by randomly sampling the structure of the MIS data in terms of the frequency of methods used, how different methods were used in combination with each other, effort, habitat, and property size. We then simulated population dynamics and removals across a range of pig densities and parameter values (Supplementary Information 1.3). Each property was simulated for 40 primary periods with six spin-up periods (periods without management). At the beginning of the spin-up period, the starting density was one of 0.3, 1.475, 2.65, 3.825, or 5 pigs/km^2^, which are equally spaced intervals between 0.3 and 5, encompassing realistic density estimates of wild pigs across North America (Davis et al., 2017; Hanson et al., 2008; Keiter et al., 2017).

#### 2.3.1 Evaluation

We evaluated posterior density estimates using bias (pigs/km^2^), normalized bias (nBias), normalized root mean squared error (nRMSE), and root mean squared log error (RMSLE). We used normalized metrics because when density increased so did bias and RMSE, reflected by the expected increase in posterior variance. Normalization reflects a more accurate way of comparing model predictions across densities. RMSLE was used when known densities included zero, as RMSLE is defined when the normalization factor equals zero. nRMSE was used on the extant density predictions only, as nRMSE is not defined when the normalization factor equals zero. Population trajectory was estimated by the slope of known densities within a property, where decreasing properties had a slope < -0.05, increasing properties had a slope > 0.05, and stable properties had a |slope| <= 0.05. Finally, random variables were considered recovered if the known value fell within the 90% confidence interval of the marginal posterior distribution.

### 2.4 Fit to MIS Data

The initial data of 3,631 properties was reduced to 1,016 properties based on removal model requirements and the first non-zero removal event. These properties were then further filtered down to 105 by using simulation results to identify properties expected to yield precise abundance estimates (see Discussion, Supplementary Information 1.4). We reduced the MIS data to the final 105 properties for two reasons: data quality and computational efficiency. The MIS data had vast amounts of spatial and temporal misalignment, meaning properties could go years without management and thus without data. This leads to computational issues because the removal IPM is iterative, estimating density for every primary period, sampled or not. The larger the dataset, the larger the ratio of unsampled to sampled primary periods, making convergence difficult.

## 3 Results

### 3.1 Simulation experiments

Across simulations and starting densities all parameters were recovered in at least 91.7% of simulations (n = 447) (Table S5), except for *ω*_*k*_. *ω*_*k*_, which estimates the percentage of unique area searched with additional snares and traps deployed, was recovered by 77.9% and 85.9% of simulations for snares and traps, respectively.

Known abundances were well recovered. When pigs were known to exist, the range in recovery rates was between 87.5-89.9% for the 0.3 and 3.825 pigs/km^2^ starting density scenarios, respectively. When the known number of pigs fell to zero, abundance was recovered 100% of the time (Table S5). Additionally, densities were recovered regardless of population growth (Fig. 2), where 90.0%, 87.5%, and 89.9% of known densities were recovered when the population was decreasing, increasing, or stable, respectively.

**Figure 2.**
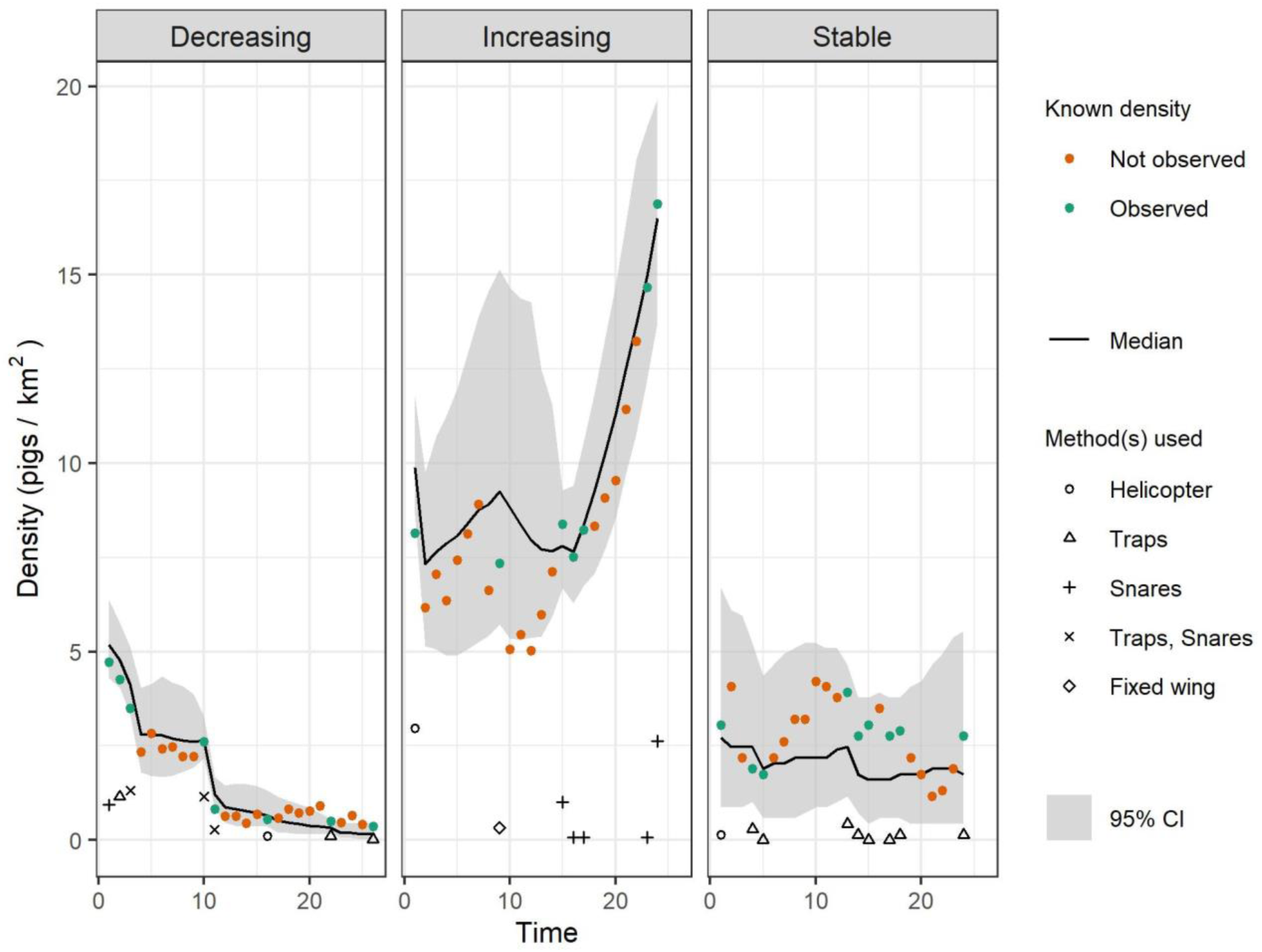
Simulated property time series for example properties with declining, increasing, and stable populations. Simulated known densities are the green and orange points where the y-axis coordinate is the known density in that primary period, and color corresponds to if removals occurred in that primary period or not. The gray ribbon and black line are posterior density estimates. The shaped points correspond to the method(s) used for removals and the y-axis coordinate for the shape is the total number of pigs removed in that primary period expressed as a density.

#### 3.1.1 How effort and method affect density estimation

The model performed best when true density was greater than 2 pigs/km^2^ and helicopters, snares, or traps were used, regardless of effort (Fig. 3). Here, the range in nBias was between -0.02 (snares, 80-100% effort, high density) and 0.05 (snares, 80-100% effort, medium density). The range in nRMSE was between 0.19 (snares, 80-100% effort, high density) and 0.39 (snares, 0-20% effort, medium density).

**Figure 3.**
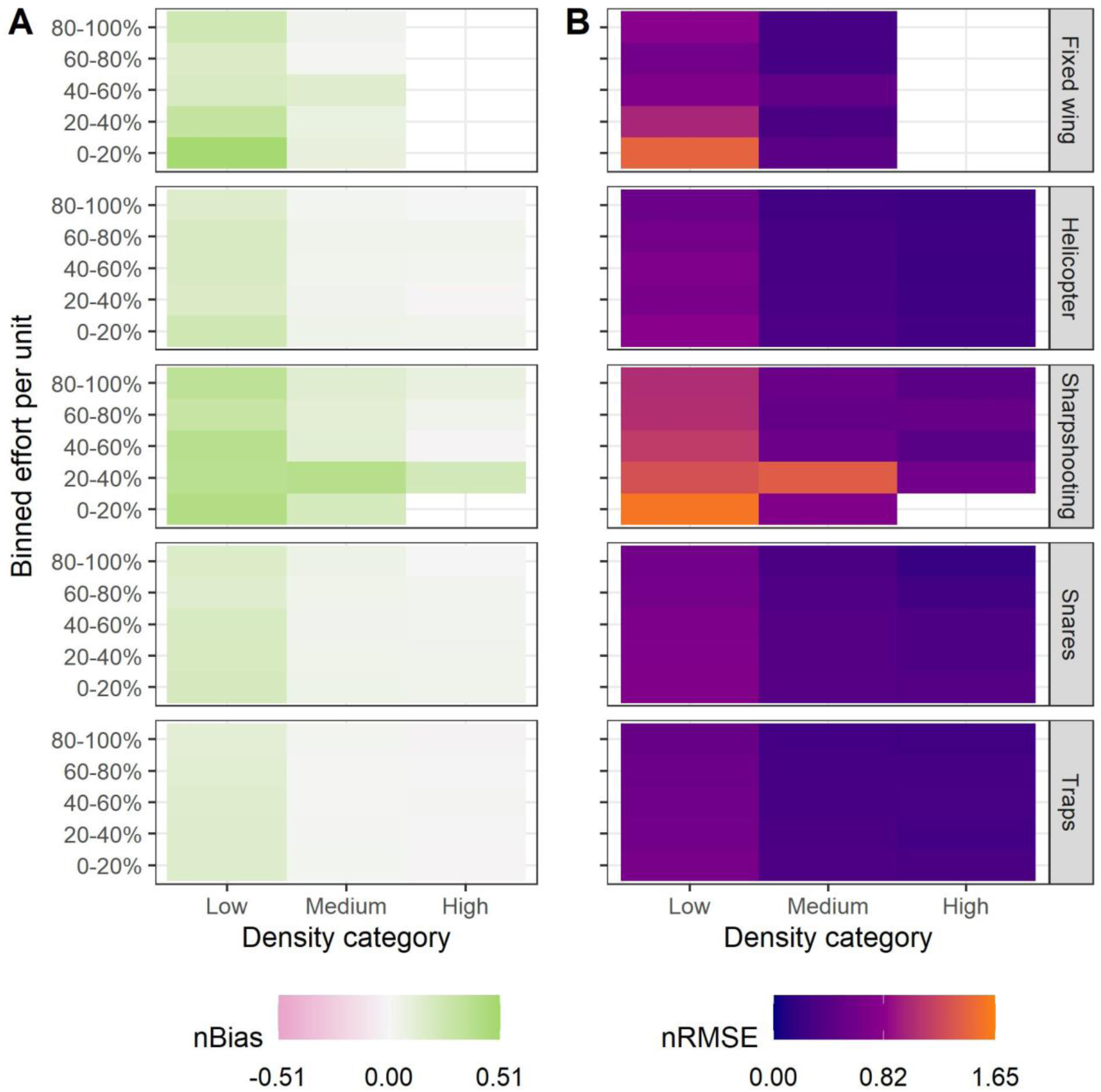
Heat maps showing how effort and method affect density estimation from the simulation study. Effort was binned into five percentile categories with respect to each method and density was binned into categories when the simulated density between 0-2 pigs/km^2^ (Low), between 2-6 pigs/km^2^ (Medium), or over 6 pigs/km^2^ (High), as per (Lewis et al., 2019). The color representation is equal to the mean error metric in each effort-density-method group. Empty spaces indicate the minimum sample size of 30 was not met for that group. A) Normalized bias and B) normalized root mean square error (RMSE).

The model made the least accurate and least precise density predictions when sharpshooting or fixed wing aircraft were used, relative effort was between 0-20%, and true density was below 2 pigs/km^2^. This scenario resulted in density predictions with a nBias of 0.48 and 0.42 and an nRMSE of 1.43 and 1.57 for fixed wing aircraft and sharpshooting, respectively. In general, as density and effort decreased, both nBias and nRMSE increased, and predictions were biased high.

#### 3.1.2 How the model performs at different levels of density

Bias was centered on zero when observations were made and that was constant across densities. Additionally, the variation in bias increased with density whether observations were made or not (Fig. 4A). Specifically, when known density was low, the 90% CI in bias was between (-0.33-0.46) and (-0.57-0.95) for observed and unobserved primary periods, respectively. When density was high the 90% CI in bias was between (-3.03-2.58) and (-6.72-4.04) for observed and unobserved primary periods, respectively. For primary periods that were not observed, density predictions tended to be biased low as density increased (Fig. 4A). Additionally, nBias was highest when density was lowest, regardless of method (Fig. 3A).

**Figure 4.**
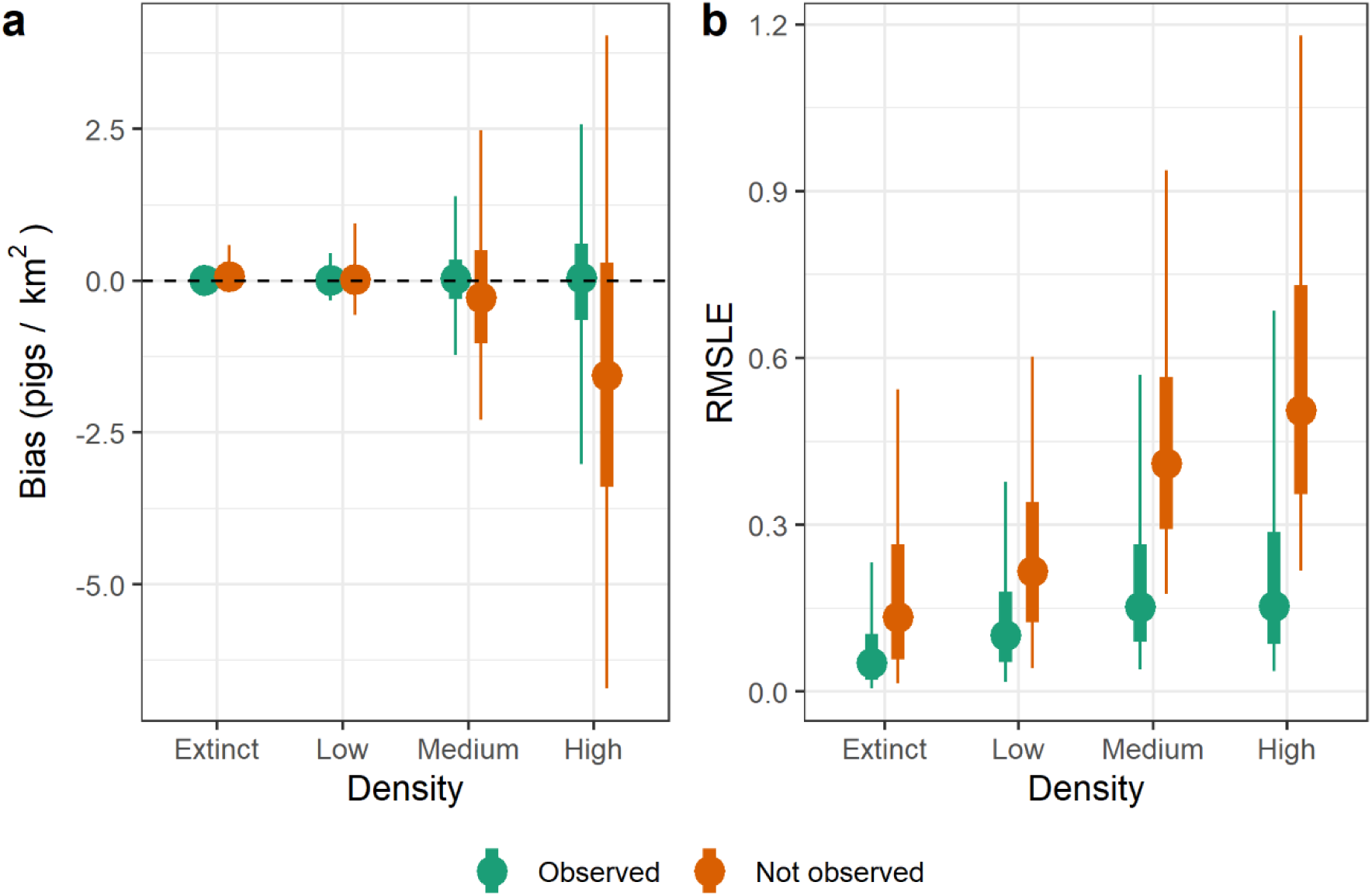
90% credible interval (CI, narrow whisker), 50% CI (wide whisker) and median (point) for the distribution of A) posterior mean bias and B) posterior root mean squared log error from the simulation study. Density categories are when the simulated density is equal to zero (Extinct), between 0-2 pigs/km^2^ (Low), between 2-6 pigs/km^2^ (Medium), or over 6 pigs/km^2^ (High), as per (Lewis et al., 2019).

RMSLE increased as density increased for unobserved primary periods (Fig. 4B). For observed primary periods where pigs were known to be alive, RMSLE was not drastically different across density categories.

### 3.2 Case study - wild pig removal data

Density estimates had a median of 1.7 pigs/km^2^ (90% CI: 0-16.3) across the 797 individual primary periods with removal efforts. Removals occurred on 105 properties from January 2014 to June 2024. Population growth, estimated as the slope across median density estimates within each property, revealed that 85 properties had decreasing wild pig populations, 12 were increasing, and eight were stable (Fig. 5). All marginal posterior distribution summaries from the fit to MIS data are reported in Table S3 with associated priors in Table S4.

**Figure 5.**
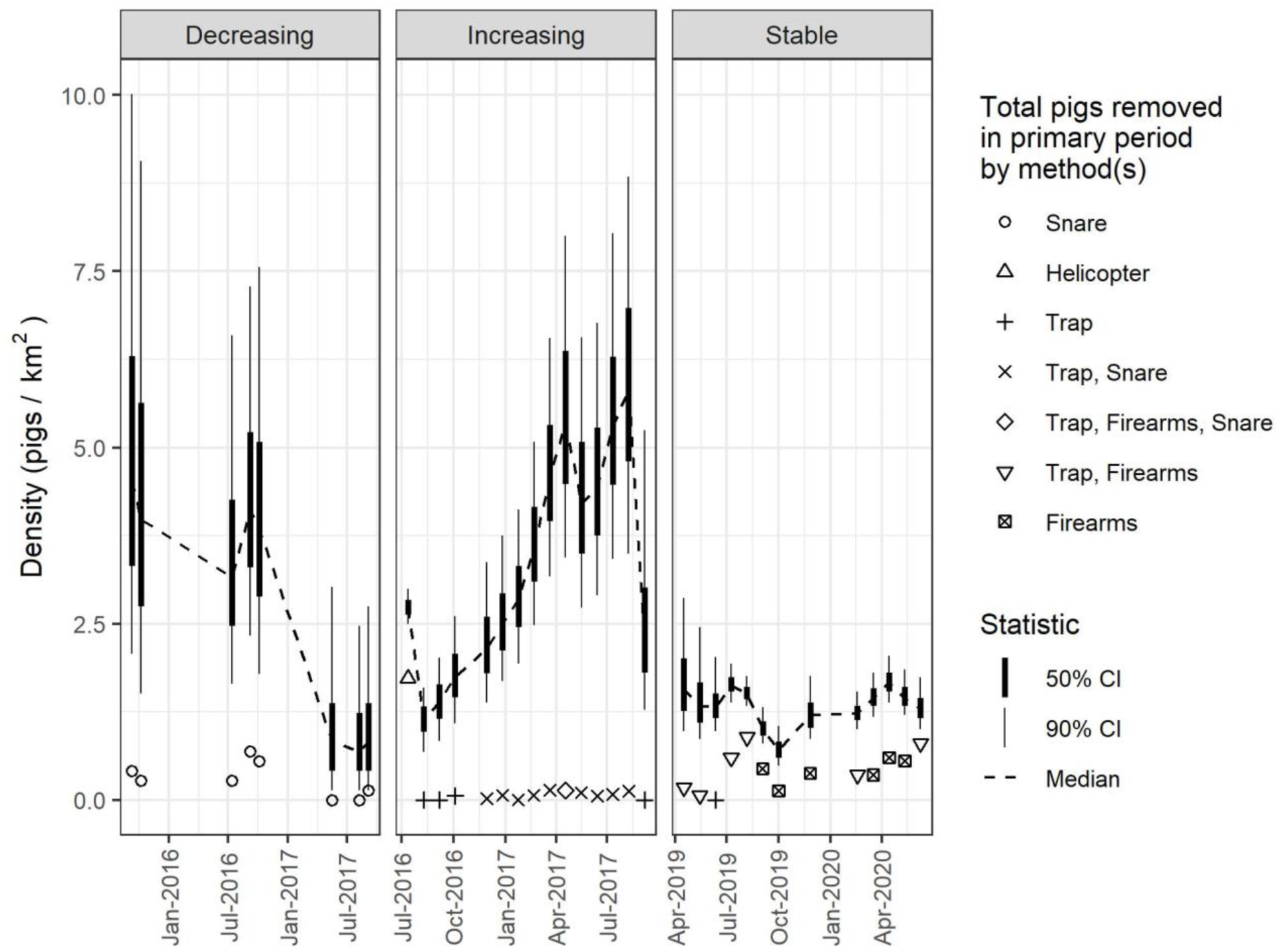
MIS property time series for three properties, one with declining, increasing, and stable populations. Posterior density estimates are represented by the dotted line and whiskers. The shaped points correspond to the method(s) used for removals and the y-axis coordinate for the shape is the total number of pigs removed in that primary period expressed as a density. The decreasing property is a 7.28 km^2^ property in Jim Wells county, TX. The increasing property is a 86.2 km^2^ property in Burnet county, TX. The stable property is a 44.9 km^2^ property in St. Landry parish, LA.

#### 3.2.1 Process model

Survival (*ϕ*_*μ*_, eqn. S1) had a median of 0.64 (90% CI: 0.61-0.66) over a 28-day period, with shrinkage (𝜓, eqn. S2) estimated at 0.66 (90% CI: 0.59-0.74). The per capita birth rate (𝜁, eqn. S12) had a median of 0.63 (90% CI: 0.59-0.68) for a 28-day period.

#### 3.2.2 Data model

Helicopters had the highest capture rate (*β*_1_) with a median of 0.98 (90% CI: 0.94-0.99), followed by traps (0.75; 90% CI: 0.56-0.88), snares (0.57; 90% CI: 0.32-0.79), sharpshooting (0.56; 90% CI: 0.35-0.76), and fixed wing aircraft (0.38; 90% CI: 0.17-0.66) (Fig. 6B).

**Figure 6.**
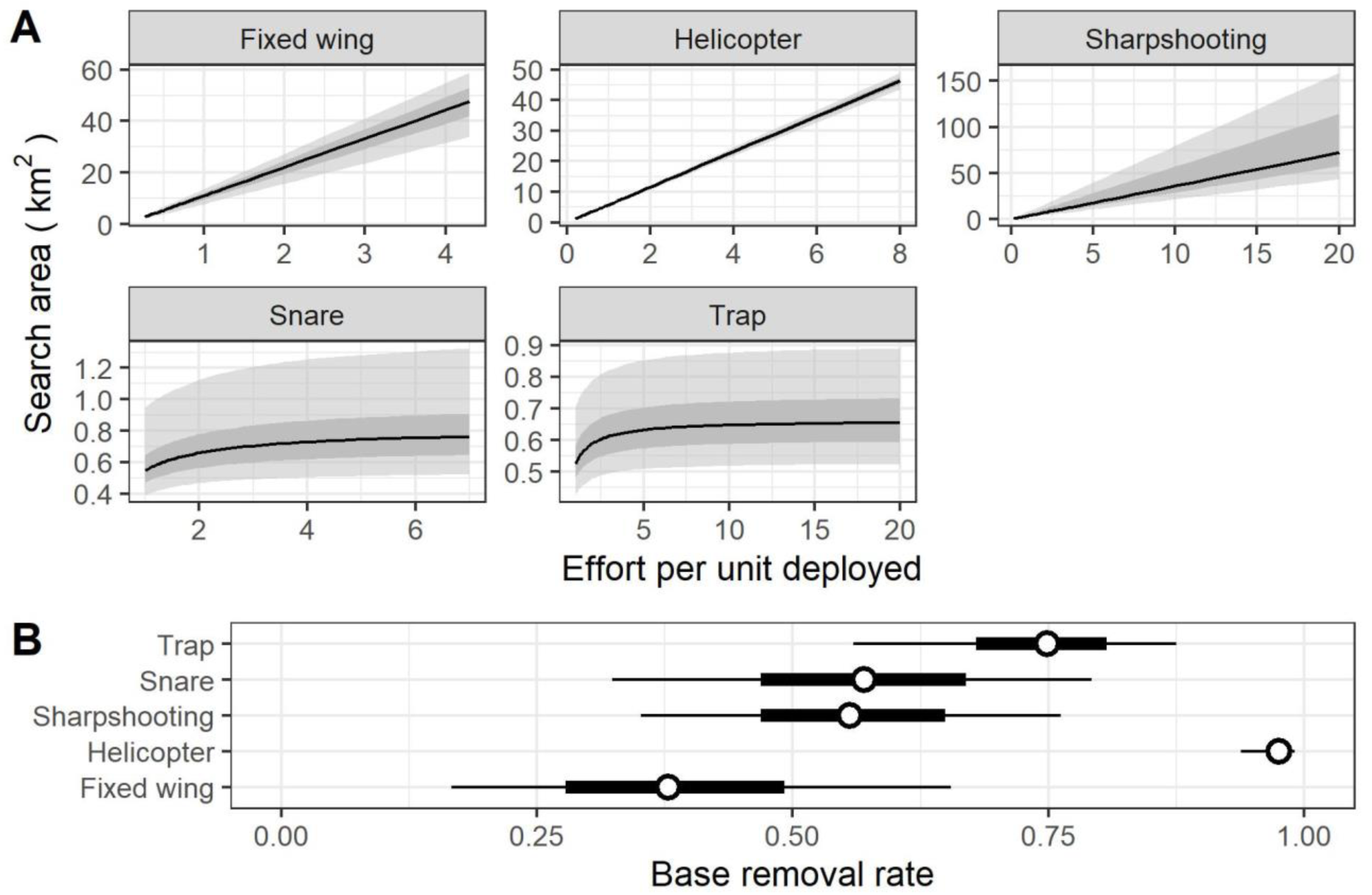
A) 90% posterior credible interval (CI; light gray area), 50% CI (dark gray area), and median (line) relating standardized effort to the potential area searched for each method. B) Posterior summary for the base removal rate (*β*_1_) for each method, the width of the narrow line is the 90% CI, the wide line is the 50% CI, and the point is the median.

The amount of area searched given effort per unit deployed increased the fastest with fixed wing aircraft (Fig. 6A), where the scaling coefficient (*ρ*_*k*_), had a median of 11.1 (90% CI: 7.89-13.7). Helicopters were next (5.81; 90% CI: 5.46-6.14), followed by hunting (3.61; 90% CI: 2.17-7.96), traps (0.45; 90% CI: 0.4-0.52), and snares (0.36; 90% CI: 0.29-0.49). The saturating effect of effort (*γ*_*k*_) was apparent for both snares and traps (Fig. 6B), with snares exhibiting a higher saturation effect with a median of 0.2 (90% CI: 0.11-0.36) and traps with a median of 0.12 (90% CI: 0.07-0.19).

The proportion of unique area (*ω*_*k*_) searched with additional traps or snares deployed was extensive, with a median of 0.05 (90% CI 0.02-0.1) for traps and 0.1 (90% CI 0.06-0.16) for snares (0 would indicate total overlap in the area searched).

Land cover differentially affected the removal rate of each method and is described in detail in the Supplementary Information 2.2.1, Fig. S6 and Table S3. But briefly, canopy cover increased the odds ratio of removal for all methods. Road density decreased the odds ratio of removal for all methods except for sharpshooting, where road density increased the odds ratio of removal. Terrain ruggedness increased the odds ratio of removal for traps, helicopters, and sharpshooting, while the removal rate of snares and fixed wing aircraft were unaffected by terrain ruggedness.

## 4 Discussion

Here, we extended a previously calibrated removal model by integrating it into an IPM framework. This approach allowed us to estimate population demographics and the search area for the five most common methods of wild pig management. This approach could be used to estimate abundance at broad scales for multiple systems, including invasive and/or game species, as removal models can inform decision-making processes and evaluate control strategies (Barron et al., 2011; Davis et al., 2016, 2017, 2022; Ramsey et al., 2009; St. Clair et al., 2013).

For invasive species which are removed using several approaches (e.g. archery, rifle, etc.), the estimation of the search area could also be informative for managers when developing regulations or hunting season length. Furthermore, detecting a reduction in survival, for example, is important for management and indicates some potential issues within a population. Population demographics could then be used to predict the long-term impacts of management actions for evaluation and refinement (Bryce et al., 2011; Gerber & Kendall, 2018; Jareb et al., 2024; Varley & Boyce, 2006).

### 4.1 How effort and method affect density estimation

Helicopters, snares, and traps affect density estimation similarly (Fig. 3), and we attribute this to how effort was recorded for helicopters, and the sample sizes for snares and traps. Flight time for helicopters was recorded with a Hobbs meter, which measures the length of time the motor is running, including travel to and from the property. This could vary significantly depending on how far the helicopter/plane has to travel. However, the Hobbs meter is the most accurate measure of effort across removal methods.

Furthermore, the area searched per hour flown is a theoretical maximum, (area searched cannot exceed the area of the property), meaning that for smaller properties it does not take many flight hours to search the entire property. Additionally, the number of animals removed by helicopters was substantial, and the number of times zero pigs were removed was minimal, which has been shown to reduce bias in density estimates (Dorazio et al., 2005).

Snares and traps were the two most used methods in the simulation study (and MIS data). Their large sample sizes likely contributed to their relatively low-biased density estimates at medium and high densities (Zhou et al., 2014).

Sharpshooting and fixed wing aircraft had the largest gradient with respect to model performance, effort, and density, where nBias and nRMSE decreased as effort and density increased (Fig. 3). The effort data for sharpshooting was tenuous, as the only values recorded were the number of sharpshooters and the number of hours they spent shooting. We did not have information about when sharpshooters employed bait, which is commonly used by managers (Davis et al., 2017; Parkes et al., 2010; Saunders et al., 1993). Additionally, sharpshooting had the fewest zero take events because sharpshooting is often targeted (baiting until there is consistent visitation at a specific site, increasing success) or opportunistic (managers are not specifically looking for wild pigs but see one while performing other duties and remove the individual). These practices also make it difficult to accurately quantify effort. If shooter ID was known, it could be used to control for effort (i.e. shooter ability) as done in large-scale monitoring programs such as the Breeding Bird Survey (Link & Sauer, 1998), and remains an area for future model development.

### 4.2 How the model performs at different levels of density

The model had the most nBias and nRMSE when density was low, which follows previous removal modeling efforts (Keiter et al., 2017). However, absolute bias was low when density was low, with 90% of simulations being within 0.7 pigs/km^2^ of known density. Ecologically, the difference of less than 1 pig/km^2^ is minimal. However, the management implications of this error could be significant, as density estimates could be used to allocate resources to remove individuals. Across the range of property sizes in the MIS data, 0.7 pigs/km^2^ translates to a possible bias of 3-104 individuals for the smallest and largest properties, respectively. And, as the mean number of pigs removed per event is less than one for the most used methods (snares and traps), removal efforts could have less impact on population reduction than intended.

### 4.3 Data criteria

The simulated properties with the most confident density estimates revealed that properties should be less than 405 km^2^ in area, *and* at least 13 pigs removed, *and* at least 24% of the primary periods have removal events, *and* at least seven removal events at a property (Table S2). This suggests that, with respect to confident density estimates, removal events need to be consistent with enough effort to thoroughly search a property, and that effort must result in multiple individuals removed. Large properties with inconsistent effort lead to inaccurate density estimates because they do not search enough relative area to find and remove individuals. Indeed, Davis et al. (2022), found that at least 30% of a property must be searched to accurately estimate density.

### 4.4 Case study - wild pig removal data

Our density estimates align with others (Keiter et al., 2017; Lewis et al., 2017, 2019; Snow et al., 2024, Taylor et al. *In review*), and we estimated a reduction in wild pig density at a majority of the properties considered. We emphasize that our technique uses only management data; we don’t use camera data or data on pig movements.

### 4.5 Catch efficiency for each removal method

Helicopters were the most efficient method (Fig. 6B) which aligns with previous work (Davis et al., 2017; Snow et al., 2024). However, aerial operations can only be used efficiently in limited conditions (low vegetation cover (Davis et al., 2018), low wind, high density populations), and are thus only applied where they are highly efficient. In contrast, other methods are applied in a wider variety of conditions (dense vegetation, low wild pig densities), where removal is known to be inefficient by any method. Thus, the efficiency estimates by removal method are biased by the conditions in which they are applied.

Traps had the second highest removal rate after helicopters. However, the area searched plays a key role in our model, and these two concepts (capture rate and area searched) must be interpreted together. For example, traps may have had a median removal rate of 0.75 *in isolation*. I.e. a pig has a 75% chance of being removed only if it is *within* the 0.6 km^2^ search area of a trap.

Our estimate of the area searched by traps was an order of magnitude less than previously estimated (Davis et al., 2017; McRae et al., 2020). Our approach differs in that we explicitly estimated a saturating effect of effort and the degree to which multiple traps search unique areas, and more accurately reflects, on average, the utility of these methods given effort. An important improvement on existing removal models because effort could be more precisely (i.e. not overprescribed) allocated to effectively search smaller properties.

We did not have data about fine scale habitat quality, trap location, trap type, and we didn’t track pigs’ movements with GPS. Given this lack of information, we show that additional traps deployed and/or additional trap nights did not increase the area searched by traps after ∼ 2.5 hours/trap. While wild pig movement has been shown to change given habitat quality (Clontz et al., 2021; West et al., 2009) and in response to trapping efforts (Bastille-Rousseau et al., 2021; Snow & VerCauteren, 2019), our approach integrates over habitat quality, trap location, and trap type to estimate the area searched by traps using data with the largest temporal and spatial extent that we are aware of. Additionally, we used county-level land cover characteristics to modify the capture rate *within* the searchable area, not to modify the searchable area itself.

We found that removal rates increased for all methods with percent canopy cover. This finding contradicts Davis et al. (2018), which did a controlled study to estimate differences in removal rates in two areas with similar density and different cover. However, landscape covariates at larger scales, such as county in our case, more likely reflect habitat suitability with respect to climate (Wiens, 1989), and wild pigs preferentially choose hardwood forests and dense canopy cover habitats (Clontz et al., 2021; T. S. Evans et al., 2024; Gray et al., 2022; Kramer et al., 2024). The length of our study and the number of properties considered also likely contributed to this discrepancy.

We found that road density increased the removal rate for sharpshooting, which follows previous studies that demonstrate ground shooting from roads and vehicles to be an efficient removal method (West et al., 2009). Nevertheless, pigs were less likely to be removed by the other four methods when road density was high. The relationship between wild pigs and roads is conflicting, where both positive (Kay et al., 2017; Snow et al., 2022) and negative (O’Brien et al., 2019) associations have been estimated. Pigs are thought to use roads for travel when resources are scarce (Clontz et al., 2021), but can also provide a barrier to movement (VerCauteren et al., 2019). Our results suggest that managers make efficient use of roads to find pigs, regardless of habitat.

## 5 Conclusion

In this study, we presented a generalizable modeling framework for estimating invasive species abundance using management data from multiple removal methods applied across divergent ecological contexts. The model accurately estimated density when removal efforts occurred, regardless of population trajectory. The model performed well across a wide range of densities, removal methods, and geographic contexts, making it a useful tool for abundance evaluation of invasive species. Additionally, this framework could be used to evaluate management impacts for national-scale programs (Jareb et al., 2024) to help with invasion control, especially on the invasion edge (Epanchin-Niell & Wilen, 2012).

