## Supplement for "Improving a removal model for evaluating density changes in a widespread invasive species"

### 1. Supplementary Methods

#### 1.1 Survival IPM

$$\phi_{\mu} \sim Beta(\alpha, \beta) \qquad (S1)$$

$$\psi \sim Gamma(1, 0.1) \qquad (S2)$$

$$\phi_{i,t} \sim Beta(\alpha_{\phi}, \beta_{\phi}) \qquad (S3)$$

Survival rate of pigs varied hierarchically across space and time. The global mean survival rate, $\phi_{\mu}$, was estimated using an informative beta distribution with hyper parameters $\alpha$ and $\beta$. $\psi$ represents the amount of shrinkage in survival across space and time. The global survival rate and shrinkage rate were used to estimate hyperparameters $\alpha_{\phi} = \phi_{\mu}\psi$ and $\beta_{\phi} = (1 - \phi_{\mu})\psi$, which were then used to draw the survival rate for each property and primary period, $\phi_{i,t}$. Hyperparameters $\alpha$ and $\beta$ for the informative prior on $\phi_{\mu}$ were parameterized by collating estimates from the literature, which can be found in Table S1. Survival estimates from these papers were converted to 4-week survival estimates to align with primary periods, as described in the Supplementary Methods.

**Table S1**

| **Paper ID** | **Survival** | **Uncertainty** | **State** | **Days** |
|---|---|---|---|---|
| Adkins & Harveson (2007) | 0.86 | 0.68-1 (95% CI) | Texas | 151 |
| Hanson et al. (2009) | 0.354 | 0.031 (SE) | Georgia | 852 |
| | 0.296 | 0.035 (SE) | Georgia | 852 |
| | 0.205 | 0.038 (SE) | Georgia | 852 |
| | 0.16 | 0.042 (SE) | Georgia | 852 |
| | 0.227 | 0.038 (SE) | Georgia | 852 |
| | 0.179 | 0.042 (SE) | Georgia | 852 |
| | 0.083 | 0.045 (SE) | Georgia | 852 |
| Hayes et al. (2009) | 0.808 | | Mississippi | 213 |
| | 0.414 | | Mississippi | 150 |
| | 0.517 | | Mississippi | 183 |
| (Gabor et al., 1999) | 0.88 | 0.09 (SD) | Texas | 364 |
| | 0.5 | 0.18 (SD) | Texas | 365 |
| | 0.4 | 0.11 (SD) | Texas | 364 |
| Gaston (2008) | 0.74 | 0.454-1 (95% CI) | Alabama | 364 |
| | 0.62 | 0.309-1 (95% CI) | Alabama | 364 |
| | 0.48 | 0.224-1 (95% CI) | Alabama | 364 |
| | 0.32 | 0.135-0.977 (95% CI) | Alabama | 364 |
| | 0.52 | 0.279-1 (95% CI) | Alabama | 364 |
| | 0.46 | 0.243-0.984 (95% CI) | Alabama | 364 |
| | 0.44 | 0.227-0.984 (95% CI) | Alabama | 364 |
| | 0.23 | 0.091-0.764 (95% | Alabama | 364 |

| | | | | |
|---|---|---|---|---|
| | | CI) | | |
| Diong (1982) | 0.345 | | Hawaii | 365 |
| | 0.283 | | Hawaii | 365 |
| | 0.135 | | Hawaii | 365 |
| | 0.061 | | Hawaii | 365 |
| | 0.403 | | Hawaii | 365 |
| | 0.342 | | Hawaii | 365 |
| | 0.283 | | Hawaii | 365 |
| | 0.125 | | Hawaii | 365 |
| | 0.05 | | Hawaii | 365 |
| Singer & Ackerman (1981) | 0.39 | | Tennessee | 211 |
| | 0.46 | | Tennessee | 211 |
| | 0.76 | | Tennessee | 211 |
| | 0.7 | | Tennessee | 211 |
| Henry & Conley (1978) | 0.585 | | Tennessee | 4747 |
| | 0.421 | | Tennessee | 4747 |
| | 0.386 | | Tennessee | 4747 |
| | 0.6 | | Tennessee | 4747 |
| | 0.528 | | Tennessee | 4747 |
| | 0.42 | | Tennessee | 4747 |
| | 0.5 | | Tennessee | 4747 |
| | 0 | | Tennessee | 4747 |
| (Barrett, 1971) | 0.856 | | California | 883 |
| | 0.711 | | California | 883 |
| | 0.416 | | California | 883 |
| | 0.856 | | California | 883 |

| | | | | |
|---|---|---|---|---|
| | 0.711 | | California | 883 |
| | 0.416 | | California | 883 |

**Table S1** - Literature used to construct the informative prior on survival. Citation is the study that information was pulled from. Survival is the observed or estimated proportion of pigs surviving over the number of days in the study, multiple rows indicates multiple sites/populations in the study. Uncertainty is the reported type of error in survival estimates. Multiple replicates for each study indicate different populations that were monitored in each study.

The studies in Table S1 report the proportion of pigs that survived over the duration of the study ($\kappa$). Therefore, we calculated the mean rate of survival per 4-week period ($\mu$) for each study $i$ given how many 4-week intervals were contained in the length of each study ($npp$). For uncertainty in survival, some papers reported error as standard deviation, and others as 95% CI. For the studies that reported 95% CI, we assumed a Gaussian distribution and converted those to standard deviation and then to variance ($\sigma_i^2$). We then scaled the variance to represent 4-week variance on survival ($\sigma_{pp,i}^2$) by the square of the scaling factor between the 4-week survival of each study and the original surviving proportion. $\alpha$ and $\beta$ were calculated using the mean 4-week survival and 4-week variance estimates.

$$\phi_{pp,i} = \kappa_i^{\frac{1}{npp_i}} \qquad (S4)$$

$$\overline{\phi_{pp}} = \frac{\sum_i^N \phi_{pp,i}}{N} \qquad (S5)$$

$$\sigma_{pp,i}^2 = \sigma_i^2 \left(\frac{\phi_{pp,i}}{\kappa_i}\right)^2 \qquad (S6)$$

$$\overline{\sigma_{pp}^2} = \frac{\sum_i^N \quad \sigma_{pp,i}^2}{N} \tag{S7}$$

$$\alpha = \frac{\overline{\phi_{pp}}}{\overline{\sigma_{pp}^2}} \tag{S8}$$

$$\beta = \frac{1 - \overline{\phi_{pp}}}{\overline{\sigma_{pp}^2}} \tag{S9}$$

#### 1.2 Fecundity IPM

$$G \sim Poisson(\nu) \tag{S10}$$

$$\nu \sim exp\big(N(2,1)\big) \tag{S11}$$

$$\zeta = \nu * \frac{28}{365} \tag{S12}$$

The birth rate, $\nu$, is constrained by the data vector $G$ which is the mean fetal litter size estimates collated in Snow et al., (2020). $\zeta$, then, is the average number of pigs added to a population through births and immigration in a 28-day period.

#### 1.3 Simulation Experiment

Below, we briefly describe how we structured the pseudo-MIS data, how ecological and removal dynamics were simulated, and the initial conditions for each simulation.

##### *1.3.1 Data set creation*

Each simulation contains 40 primary periods with 110 “empty” properties, i.e. nothing has been assigned to these properties such as method(s) used for removals, the frequency at which removals happened, the area of the property, or effort. Below, we use a hypothetical 2-

method property as a working example to explain the procedure used to "fill" each property with attributes. The same procedure was used for all $n$-method properties.

First, we assigned each property a starting primary period, which was randomly chosen between 1-24, so that at a minimum at least 16 primary periods could theoretically have removal events. For our example property, the first primary period when removals occur is primary period 14. Then, we assigned each of the 110 properties the number of methods they would use (where $n = 1, 2 \ldots 5$), where the proportion of properties that were assigned 1-method was equal to the proportion of properties in the real MIS data that use a single method. The same procedure of assigning $n$-methods was done for all $n$-method properties, so that the number of properties assigned to $n$-method properties were 41, 49, 18, 4, and 1 for 1- to 5-method properties, respectively. Each property was assigned a single method for 1-method properties or a combination of methods for multiple-method properties.

For our example 2-method property, we first determined the relative frequency of all pairings of methods found in the real MIS data and used those frequencies as weights to randomly assign each 2-method property a pair of methods. Our example property will be assigned snares and traps, as these two methods were the most frequently paired set of methods, which were employed by 52.2% of 2-method properties in the real MIS data.

After we assigned a method(s) to each property, we assigned an area ($km^2$) to each property. Areas were randomly chosen from the real MIS data given the specific method combination for that property. For our example property, we pulled property areas from the real MIS data for properties that only use snares and traps, and then randomly selected one of those areas. This ensures that, for example, properties assigned to use traps or snares will have relatively smaller property areas than properties assigned to use helicopters, because helicopters are used on larger properties in the real MIS data. Our example property was randomly assigned an area of 8.09 $km^2$, which is the area of a property in the real MIS data that uses only snares and traps.

Now we have an example property of 8.09 $km^2$ that uses snares and traps, whose time series starts at primary period 14. The next step is to determine in which primary periods removals occur. This means determining in which primary periods snares and traps are used together or separately, and how many replicate events occur when a method is used. From the properties in the MIS data that only use snares and traps, we calculated the return intervals (the number of primary periods it takes for a method(s) to be used again in a property) for snares, traps, and snares and traps used together. This creates a table that has the return interval tied to a single method (snares *or* traps) or both methods (snares *and* traps). We randomly sampled the rows of this table to determine the next primary period that removals occurred, and which method(s) are used in that primary period. We repeated the sampling until the maximum primary period in the time series was less than or equal to 40.

To assign the number of replicate events by each method in a primary period, we again sampled from the properties in the MIS data that only use snares and traps by calculating the number of replicate events for each method (separately and jointly), then randomly sampled from those values. The order of each removal event was randomly assigned, regardless of method. In our example property, the first sampled primary period was assigned one snare event and five trapping events, and those were ordered as: traps, snares, traps, traps, traps, traps.

Before we simulate the removal of pigs, we need to determine the effort of each removal event, which again was determined by sampling from the real MIS data given the method. This includes the number of units (snares, traps, etc.) deployed, then we sampled the normalized effort (hours per unit deployed) spent removing given the number of units deployed. For example, in the first removal event in our example property, we found the number of traps deployed for all trapping events in the real MIS data and randomly sampled one of those values, which in our example was three traps. Then, for all trapping events that used three traps, we randomly chose a normalized effort value, which in our example is one hour spent trapping per trap.

The operating procedure described above was completed for each primary period in each of the 110 properties in each simulation and provides the structure from which ecological and removal dynamics were simulated. Each simulation was assigned a starting density, which was one of 0.3, 1.475, 2.65, 3.825, or 5 pigs/km$^2$. These values were chosen to be equally spaced between 0.3 and 5, which represents the range in estimated wild pig densities across the continental US (VerCauteren et al., 2019).

##### *1.3.2 Ecological and removal dynamics*

For each property, there was a spin-up period of six primary periods where the initial abundance at each property was the assigned starting density multiplied by the property area. After the spin-up period, the property was discarded from the simulation if the population went extinct, or became unrealistically dense, with a maximum cutoff density of 10 pigs/km$^2$. Population dynamics were simulated using the ecological model described above and in the main text. If the property wasn't discarded, the spin-up period was discarded, and the initial abundance was set to the abundance at the end of the spin-up period. We then simulated removal and ecological dynamics for each property using the data model, ecological model, and pseudo-data described above. If a primary period was assigned to have removals take place, those happened first and the remaining population was used as the initial abundance for the population model, otherwise the initial abundance was the abundance from the previous primary period. When removals occurred, we ensured that the number of pigs removed could not exceed the total population size.

##### *1.3.3 Model parameters*

For each simulation (a set of 110 properties and one of the five starting densities), model parameters for the data model were chosen at random (Table 1). The upper ends for $\rho$ for the aerial methods, snares, and traps were informed by Davis et al. (2017) and McRae et al. (2020), where we added a small buffer to the maximum area estimated for those methods. For sharpshooting, we assumed that a single hunter could search a maximum of five km$^2$ in a single bout of hunting.

For the saturating constant ($\gamma$), the distribution for traps and snares was optimized to match the values we expected based on 100,000 simulated draws of the function relating area and effort, minimizing the sum of squared errors between the prior modes and 99% prior quantiles, evaluated at two points: one corresponding to one unit of effort (e.g. one trap night), and a second corresponding to a larger effort value for which we had prior information (e.g. 14 trap nights). Our prior expectation was that area sampled by effort saturates for snares and traps, but not for aerial methods and sharp shooting.

For the ecological model, the 4-week survival rate ($\phi_\mu$) was set to 0.78 and shrinkage ($\psi$) was set to 5 for every simulation. These values were chosen because they allowed for realistic variation in simulated abundance. I.e. most properties did not grow exponentially, and some went extinct due to environmental/demographic stochasticity, while most properties had a "natural" variation in abundance through time.

##### *1.3.4 MCMC*

Our state-space IPM was fit to each simulated data set using a MCMC implemented in NIMBLE (de Valpine et al., 2022, 2017). Each simulation was run for a total of 400K iterations across three chains, with the first half of each chain discarded as burn-in. We then calculated the point scale reduction factor (PSRF) (Gelman & Rubin, 1992) for each parameter, and discarded simulations that did not reach convergence (PSRF $\leq$1.1). For some simulations we also looked at trace plots to decide convergence, especially when $\psi$ was > 1.1 but nonetheless looked converged. After removing simulations that did not converge, the number of properties remaining for analysis was 35,835, and ranged from 5,637-8,919 for the 0.3 and 3.825 starting densities, respectively.

#### 1.4 Fit to MIS Data

The initial MIS data contained 3,631 properties, which was reduced to 1,016 properties based on removal model requirements and the first non-zero removal event. These properties were then further filtered down to 105 by using simulation results to identify properties expected to yield precise abundance estimates. We reduced the MIS data to the final 105 properties for two reasons: data quality and computational efficiency. The MIS data had vast amounts of spatial and temporal misalignment, meaning properties could go years without management and thus without data. This leads to computational issues because the removal IPM is iterative, estimating density for every primary period, sampled or not. The larger the dataset, the larger the ratio of unsampled to sampled primary periods, making convergence difficult.

Ultimately, we fit the model to a subset of all available properties in the MIS data. We filtered out properties for two reasons. The first filtering occurred due to the requirements of removal models, which require at least two removal events in a primary period, and that a property have at least two primary periods. We also conditioned on the first positive capture. This winnowed the data from 3631 to 1016 properties (Table S1). We binned individual removal events into 28-day primary periods, which started on the first recorded removal event. The 28-day interval was chosen because it is long enough to keep most properties with sparse removal data, and short enough that the assumption of population closure within a primary period was reasonable.

For the second filtering step, we used the simulation results to filter the real MIS properties down to a set that we expected to yield precise abundance estimates. For each property in the simulation experiment, we calculated mean percentage error (MPE pigs/km$^2$), normalized root mean squared error (NRMSE), and mean bias (pigs/km$^2$) with respect to posterior density vs known density, and then filtered to properties that had less than 25% MPE, less than 0.5 NRMSE, and between -0.5 and 0.5 mean bias. 20.4% (n=7306) of simulated properties fit these criteria. From this sample of simulated properties, we calculated the 5% and 95% quantiles of simulated property characteristics (e.g. property area, total number of pigs removed, total number of removal events, etc.), where these quantiles provided the range of property characteristics that were used to winnow the MIS data. For example, the 7306 simulated properties that fit our selection criteria had property areas with a 90% CI of 4.05-405 km$^2$. Therefore, the properties in the MIS data that we fit our model to also had to have an area within this range. A complete description of the property characteristics that were used to filter the MIS data are provided in Table S2. This winnowing brought the final number of properties to 105.

Within primary periods, when the order of removal events was ambiguous, semi-stochastic ordering was applied based on expert knowledge (informal discussions with Wildlife Services feral swine management specialists and site visits) of how removals are typically conducted. When multiple removal methods were used and the order of these events could not be determined, we assumed that traps and snares occurred first, then helicopters and fixed wing aircraft, and finally sharp shooting.

##### *1.4.1 MCMC*

Our state-space IPM was fitted to the 105 MIS properties using a MCMC implemented in NIMBLE (de Valpine et al., 2022, 2017). We manually changed the sampler for $\beta_{1,1-K}$ (intercept, eqn. 6) to a random walk blocked across the $K$ methods with an adaptive interval set to 100 and a rate of decay for the scale adaption factor set to 0.6. We also changed the sampler for $\nu$ (eqn. S11) to the default slice sampler. We ran seven chains in parallel until all parameters reached an effective sample size (ESS) of 1000 and a PSRF < 1.1. We ran a total of 651K iterations, burn-in occurred at iteration 18,237 with the extra iterations to achieve the desired ESS.

#### 1.5 MODIS data

The moderate-resolution imaging spectroradiometer (MODIS) vegetation continuous field data was used to describe percent tree cover at a 250-meter resolution (DiMiceli et al., 2015). The average percent tree cover was calculated for each county. Terrain ruggedness was calculated using Riley et al. (1999) terrain ruggedness index using 30-meter resolution digital elevation models collected by the Shuttle Radar Topography Mission (SRTM) (Jpl, 2013). Calculation of terrain ruggedness was done using the R spatialEco package (Evans & Murphy, 2023). The average terrain ruggedness index was calculated for each county. Road density was calculated using U.S. Census Bureau TIGER Line data that contains all known U.S. road infrastructure. TIGER road infrastructure data was acquired using the R tigris package (Walker, 2016). Road density was calculated by summing the total linear length of all roads (km) in a county and dividing by the area of the county. Percent canopy cover, road density, and average terrain ruggedness were centered and scaled to one standard deviation.

### 2. Supplementary Results

#### 2.1 Simulation Results

The parameters that estimate capture rate ($\beta_k$, eqn. 6), were well recovered. The median residuals for each parameter were centered on zero. Sharpshooting and fixed wing aircraft had the widest distribution in residuals compared to other methods (Figure S1). Nonetheless, greater than 90% of simulations recovered all $\beta_k$.

The parameter that estimates the saturation constant for snares and traps, $\gamma_k$ (eqn. 5), was well recovered, especially for traps, and there was a slight negative bias in residuals for snares (Figure S2A). On average across all simulations, the trend in median posterior values matched the trend in known parameter values (Figure S2B).

The parameter that scales the searchable area by method, $\rho_k$ (eqns. 4 & 5), was well recovered with median residuals centered on zero for all methods (Figure S3A). Estimates of $\rho_k$ were most confident for helicopters and traps. On average, the trend in median posterior values across simulations matched the trend in known values for all methods except for snares, where there was a slight positive bias at the upper end of known values (Figure S3B).

The parameter that estimates the percentage of unique area searched with additional units for snares and traps, $\omega_k$, had residuals centered on zero (Figure S4A). The trend in posterior median estimates matched the trend in known values better for traps than snares, and these trends tended to improve as simulation starting density increased (Figure S4B).

Demographic parameters were recovered by nearly all simulations and starting densities (Table S4). Residuals were centered on zero and extremely confident for survival ($\phi_\mu$) and mean litter size ($\nu$) (Figures S5A & B). Median residuals for the shrinkage parameter on survival ($\psi_\phi$) had a slight positive bias, and the distribution of residuals became more confident as starting density increased in the simulations (Figure S5C). Nonetheless, 97.54% of simulations recovered $\psi_\phi$ (Table S2).

#### 2.2 Fit to MIS data

##### *2.2.1 The effect of land cover on removal rates*

Canopy cover increased the odds ratio of removing pigs for all methods. For a one standard deviation increase in percent canopy cover, the odds ratio of removing a wild pig with helicopters increased by a factor of 21.1, with sharpshooting by 18.5, with snares by 5.7, with fixed wing aircraft by 5.3, and with traps by 2.5.

A one standard deviation decrease in road density decreased the odds ratio of removing a wild pig with helicopters by a factor of 19.1, with snares by 8.7, with traps by 5.4, and with fixed wing aircraft by 4.3. A one standard deviation increase in road density increased the odds ratio of removing a wild pig with sharpshooting by a factor of 7.7.

A one standard deviation increase in terrain ruggedness increased the odds ratio of removing a wild pig with traps by a factor of 6.7, with sharpshooting by 4.1, with helicopters by 2.4, and with snares by 1.2. A one standard deviation decrease in terrain ruggedness decreased the odds ratio of removing a pig with fixed wing aircraft by a factor of 1.2.

#### 3. Supplementary Figures

**Figure S1**

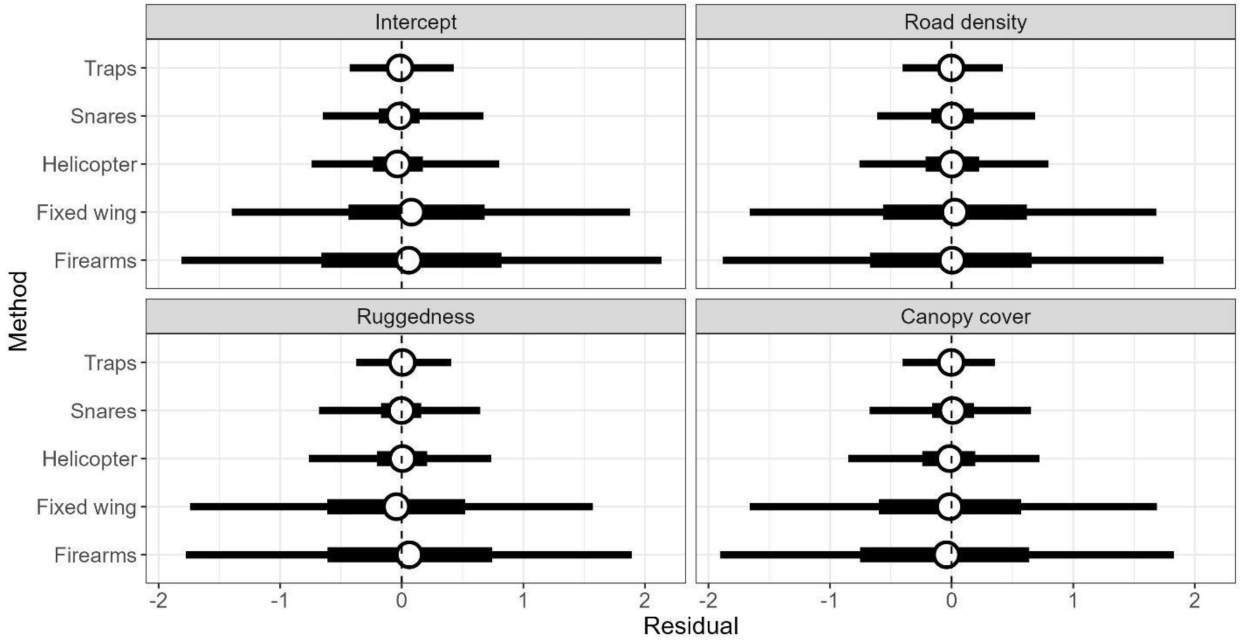


Figure S1: Residual posterior summaries for each coefficient that estimates capture rate across all simulations. Whisker plots show the median (circle), 50% confidence interval (wide line), and 90% confidence interval (narrow line).

**Figure S2**

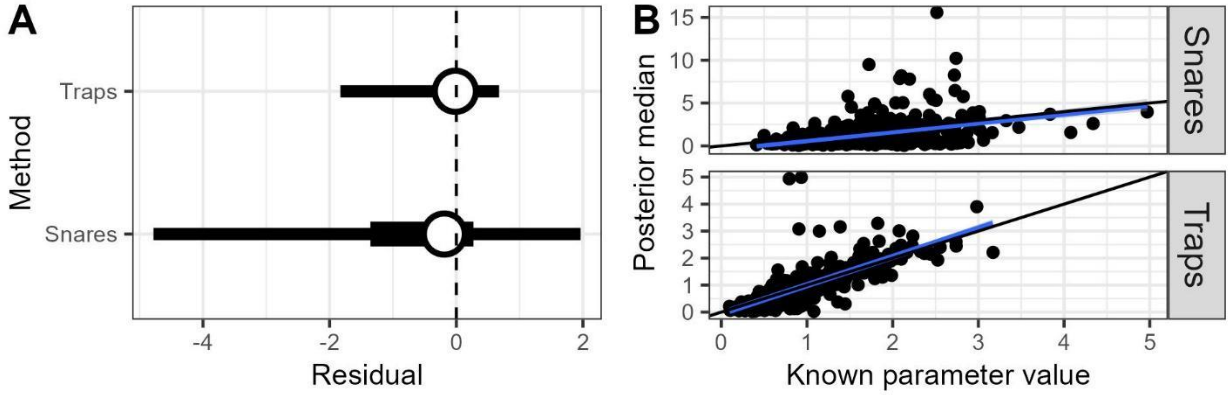


Figure S2: A) Residual posterior summaries for the saturation constant estimating area searched for snares and traps. Whisker plots show the median (circle), 50% confidence interval (wide line), and 90% confidence interval (narrow line). B) Accuracy (posterior median) of posterior estimates vs known parameter values for the saturation constant estimating area searched for snares and traps. The blue line is the best fit linear trend in posterior median values, and the black line is the 1:1 line.

**Figure S3**

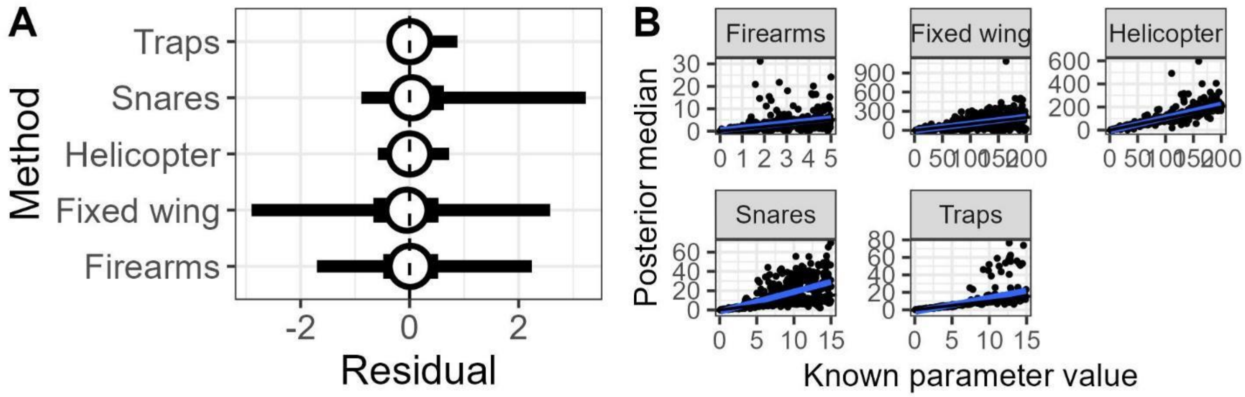


Figure S3: A) Residual posterior summaries for the scaling factor between effort and area searched. Whisker plots show the median (circle), 50% confidence interval (wide line), and 90% confidence interval (narrow line). B) Accuracy (posterior median) of posterior estimates vs known parameter values for the scaling factor between effort and area searched. The blue line is the best fit linear trend in posterior median values, and the black line is the 1:1 line.

**Figure S4**

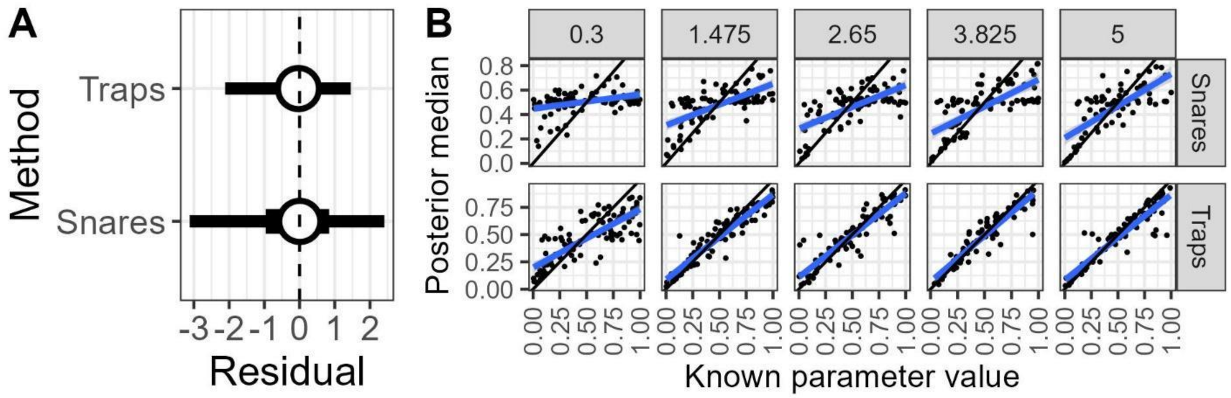


Figure S4: A) Residual posterior summaries for the percentage of unique area searched with additional units. Whisker plots show the median (circle), 50% confidence interval (wide line), and 90% confidence interval (narrow line). B) Accuracy (posterior median) of posterior estimates vs known parameter values for percentage of unique area searched with additional units; facet columns represent the simulation starting density. The blue line is the best fit linear trend in posterior median values, and the black line is the 1:1 line.

**Figure S5**

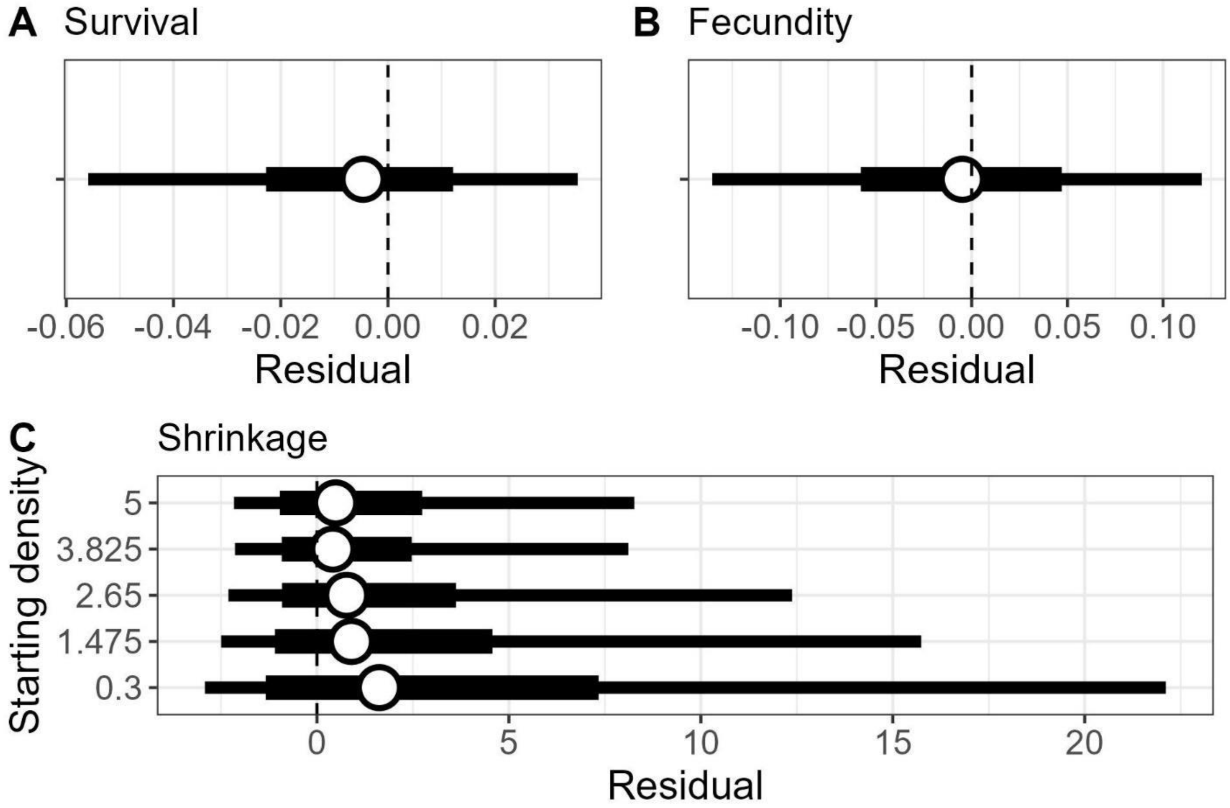


Figure S5: Residual posterior summaries for A) survival, B) per-capita recruitment, and C) shrinkage for the hierarchical survival model. Whisker plots show the median (circle), 50% confidence interval (wide line), and 90% confidence interval (narrow line).

**Figure S6**

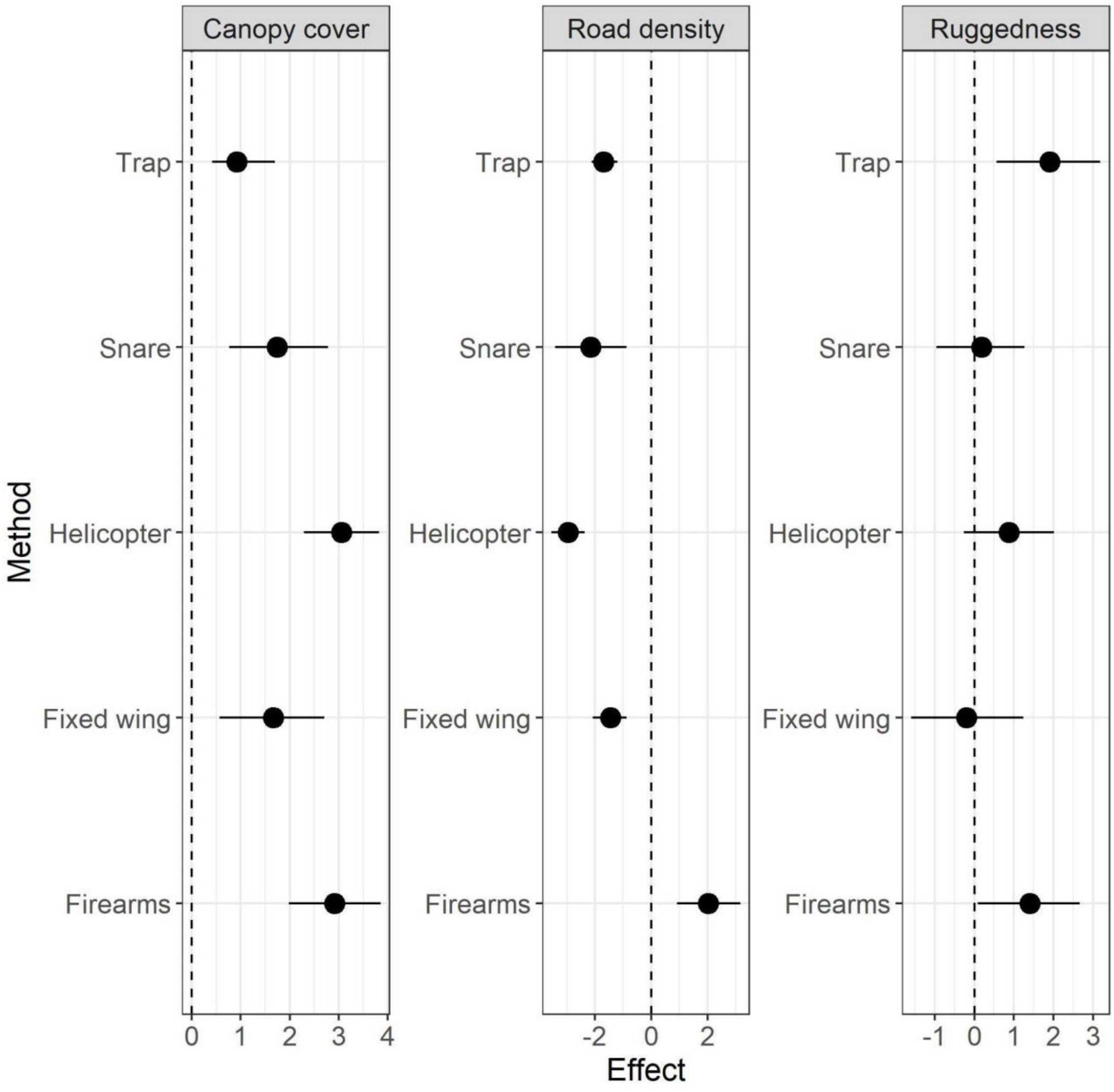


Figure S6: Poster summaries for the effect of county-level landscape covariates on removal rate by method. Median (point) and 90% credible interval (whisker) displayed.

#### 4. Supplementary Tables

**Table S1**

| | All data | | Removal model | | Fit to data | |
|---|---|---|---|---|---|---|
| State | n county | n property | n county | n property | n county | n property |
| ALABAMA | 13 | 58 | 4 | 6 | NA | NA |
| ARKANSAS | 38 | 149 | 10 | 15 | NA | NA |
| FLORIDA | 19 | 50 | 6 | 18 | 2 | 4 |
| GEORGIA | 6 | 28 | 5 | 15 | 1 | 1 |
| INDIANA | 1 | 1 | NA | NA | NA | NA |
| KANSAS | 1 | 1 | NA | NA | NA | NA |
| LOUISIANA | 38 | 194 | 10 | 26 | 1 | 1 |
| MISSISSIPPI | 24 | 109 | 12 | 39 | 3 | 4 |
| MISSOURI | 24 | 82 | 6 | 8 | NA | NA |
| NEW MEXICO | 6 | 11 | 3 | 3 | NA | NA |
| NEW YORK | 1 | 1 | NA | NA | NA | NA |
| NORTH CAROLINA | 25 | 56 | 3 | 3 | 1 | 1 |
| OHIO | 2 | 2 | NA | NA | NA | NA |
| OKLAHOMA | 15 | 246 | 12 | 61 | 6 | 14 |
| PENNSYLVANIA | 3 | 3 | NA | NA | NA | NA |
| SOUTH CAROLINA | 19 | 57 | 8 | 12 | 1 | 1 |
| TENNESSEE | 6 | 6 | NA | NA | NA | NA |
| TEXAS | 169 | 2564 | 121 | 808 | 45 | 79 |
| VIRGINIA | 6 | 9 | 1 | 1 | NA | NA |
| TOTAL | 420 | 3631 | 202 | 1016 | 60 | 105 |

Table S1: Data summary by state across data sets, n county and n property are the total number of counties and properties within each state. “All data” columns refer to all the MIS data before any winnowing. “Removal model” columns represent the subset of all data that could theoretically be applied to a removal model. “Fit to data” columns represent the ideal subset of properties to be fit with our model that were identified after the simulation study.

**Table S2**

| Property characteristic | Minimum | Maximum |
|---|---|---|
| Property area ($km^2$) | 4.05 | 405 |
| Total number of pigs removed | 13 | 858 |
| Total number of removal events | 7 | 117 |
| Total number of observed primary periods | 2 | 18 |
| Total time series length (number of primary periods) | 3 | 34 |
| Proportion of primary periods observed | 0.24 | 1 |
| Total effort | 13.9 | 263 |
| Total units deployed | 10 | 653 |

Table S2: Property characteristics identified from the simulation study that were used to winnow the MIS data into the final data set used to fit the Bayesian model. Total effort is the sum number of hours spent removing pigs per unit deployed regardless of method. Total units deployed is the total number of units deployed at a property regardless of method.

**Table S3**

| Parameter | Median | 5% | 95% | Mean | SD |
|---|---|---|---|---|---|
| $\beta_{1,sharpshooting}$ | 0.22 | -0.61 | 1.16 | 0.25 | 0.54 |
| $\beta_{1,fixed\ wing}$ | -0.50 | -1.61 | 0.64 | -0.49 | 0.68 |
| $\beta_{1,helicopter}$ | 3.68 | 2.72 | 4.68 | 3.69 | 0.59 |
| $\beta_{1,snare}$ | 0.28 | -0.74 | 1.34 | 0.29 | 0.64 |
| $\beta_{1,trap}$ | 1.09 | 0.24 | 1.95 | 1.09 | 0.51 |
| $\beta_{road\ density,sharpshooting}$ | 2.04 | 0.91 | 3.17 | 2.03 | 0.69 |
| $\beta_{ruggedness,sharpshooting}$ | 1.40 | 0.08 | 2.67 | 1.39 | 0.79 |
| $\beta_{canopy\ cover,sharpshooting}$ | 2.92 | 1.98 | 3.85 | 2.92 | 0.57 |
| $\beta_{road\ density,fixed\ wing}$ | -1.45 | -2.08 | -0.87 | -1.45 | 0.37 |
| $\beta_{ruggedness,fixed\ wing}$ | -0.19 | -1.59 | 1.23 | -0.18 | 0.84 |
| $\beta_{canopy\ cover,fixed\ wing}$ | 1.67 | 0.57 | 2.71 | 1.65 | 0.65 |
| $\beta_{road\ density,helicopter}$ | -2.95 | -3.55 | -2.36 | -2.95 | 0.36 |
| $\beta_{ruggedness,helicopter}$ | 0.87 | -0.27 | 2.02 | 0.86 | 0.70 |
| $\beta_{canopy\ cover,helicopter}$ | 3.05 | 2.29 | 3.82 | 3.05 | 0.47 |
| $\beta_{road\ density,snare}$ | -2.16 | -3.41 | -0.87 | -2.16 | 0.77 |
| $\beta_{ruggedness,snare}$ | 0.18 | -0.95 | 1.27 | 0.17 | 0.68 |
| $\beta_{canopy\ cover,snare}$ | 1.74 | 0.76 | 2.78 | 1.75 | 0.61 |
| $\beta_{road\ density,trap}$ | -1.69 | -2.11 | -1.20 | -1.68 | 0.27 |
| $\beta_{ruggedness,trap}$ | 1.90 | 0.56 | 3.18 | 1.89 | 0.79 |
| $\beta_{canopy\ cover,trap}$ | 0.92 | 0.41 | 1.69 | 0.97 | 0.39 |
| $\gamma_{snare}$ | 0.20 | 0.11 | 0.36 | 0.22 | 0.08 |
| $\gamma_{trap}$ | 0.12 | 0.07 | 0.19 | 0.12 | 0.04 |
| $\rho_{sharpshooting}$ | 3.61 | 2.17 | 7.96 | 4.31 | 1.88 |
| $\rho_{fixed\ wing}$ | 11.10 | 7.89 | 13.69 | 10.99 | 1.80 |
| $\rho_{helicopter}$ | 5.81 | 5.46 | 6.14 | 5.81 | 0.21 |
| $\rho_{snare}$ | 0.36 | 0.29 | 0.49 | 0.38 | 0.08 |
| $\rho_{trap}$ | 0.45 | 0.40 | 0.52 | 0.45 | 0.04 |
| $\omega_{snare}$ | 0.10 | 0.06 | 0.16 | 0.11 | 0.03 |
| $\omega_{trap}$ | 0.05 | 0.02 | 0.10 | 0.06 | 0.02 |
| $\nu$ | 8.27 | 7.72 | 8.83 | 8.27 | 0.34 |
| $\phi_{\mu}$ | 0.64 | 0.61 | 0.66 | 0.64 | 0.02 |
| $\psi_{\phi}$ | 0.66 | 0.59 | 0.74 | 0.66 | 0.05 |

Table S3: Marginal posterior distribution summaries for all parameters in the model after fitting to the MIS data.

**Table S4**

| **Parameter** | **Prior** |
|---|---|
| $\beta_{1-4,1-K}$ | $Gaussian(0, \tau = 1)$ |
| $\gamma_{1-2}$ | $exp\left(Gaussian(0, \tau = 3)\right)$ |
| $\rho_1$ | $exp\left(Gaussian(0, \tau = 2)\right)$ |
| $\rho_2$ | $exp\left(Gaussian(0, \tau = 1)\right)$ |
| $\rho_3$ | $exp\left(Gaussian(0, \tau = 1)\right)$ |
| $\rho_4$ | $exp\left(Gaussian(0, \tau = 3)\right)$ |
| $\rho_5$ | $exp\left(Gaussian(0, \tau = 3)\right)$ |
| $\omega_{1-2}$ | $\left(Gaussian(0, \tau = 1)\right)$ |
| $\nu$ | $exp\left(Gaussian(2, \tau = 1)\right)$ |
| $\phi_\mu$ | $Beta(3.2, 0.2)$ |
| $\psi_\phi$ | $Gamma(1, 0.1)$ |

Table S4: Priors for fitting to the MIS data. Gaussian distributions were parameterized using precision ($\tau$). $\gamma$, $\rho$, and $\nu$ were modeled on the log scale. The differing precision terms for $\rho$ reflect our prior belief that methods will have different uncertainty in how effort scales to area searched. The informative prior on $\phi_\mu$ was constructed via the Supplementary Methods described above from the studies in Table S1.

**Table S5**

| Parameter | sharpshooting | Fixed Wing | Helicopter | Snares | Traps |
|---|---|---|---|---|---|
| $\beta_1$ | 93.29 | 95.75 | 91.72 | 93.51 | 95.53 |
| $\beta_2$ | 94.41 | 95.30 | 95.97 | 92.84 | 92.62 |
| $\beta_3$ | 93.29 | 94.85 | 95.97 | 93.06 | 94.18 |
| $\beta_4$ | 91.95 | 94.18 | 93.96 | 95.30 | 94.41 |
| $\gamma$ | | | | 96.64 | 92.62 |
| $\rho$ | 95.75 | 94.41 | 93.74 | 95.53 | 94.63 |
| $\omega$ | | | | 77.85 | 85.91 |

Table S5: The percent of simulations that recovered the known parameter value for parameters in the data model. Recovered is defined as the known parameter value being within the central 90% confidence interval of the marginal posterior distribution for each parameter. The total number of simulations was 447. $\beta_1$ is the intercept for calculating removal rates, $\beta_{2-4}$ are coefficients tied to landscape covariates in the removal rate equation (in order 2-4: road density, terrain ruggedness, and canopy cover).

**Table S6**

| Parameter | Percent recovered |
|---|---|
| $\phi_\mu$ | 98.66 |
| $\psi_\phi$ | 97.54 |
| $\nu$ | 100 |

Table S6: The percent of simulations that recovered the known parameter value for parameters in the process model. Recovered is defined as the known parameter value being within the central 90% marginal posterior distribution for each parameter. The total number of simulations was 447.

**Table S7**

| Start density | Percent recovered (Extant) | Percent recovered (Extinct) |
|---|---|---|
| 0.3 | 87.5 | 100 |
| 1.475 | 88.5 | 100 |
| 2.65 | 89.1 | 100 |
| 3.825 | 89.9 | 100 |
| 5 | 89.7 | 100 |

Table S7: The percent of simulations that recovered the known abundance value for each simulation starting density. Recovered is defined as the known abundance value being within the central 90% marginal posterior distribution for abundance. Percent recovered is split between when pigs are known to inhabit the simulated property (extant) vs known to be extinct.
